# Dopamine D2 receptor upregulation in dorsal striatum in the *LRRK2*-R1441C rat model of early Parkinson’s disease revealed by *in vivo* PET imaging

**DOI:** 10.1101/2023.08.09.550512

**Authors:** Teresa Delgado-Goñi, Natalie Connor-Robson, Milena Cioroch, Stephen Paisey, Christopher Marshall, Emma L. Lane, David Hauton, James McCullagh, Peter J. Magill, Stephanie J. Cragg, Clare E. Mackay, Richard Wade-Martins, Johannes C. Klein

## Abstract

*LRRK2* mutations are the most common cause of dominantly inherited Parkinson’s disease (PD). Here, we conducted PET imaging in aged transgenic rats carrying human pathogenic *LRRK2* R1441C or G2019S mutations with [^18^F]FDOPA and dopamine D2/3 receptor ligand [^18^F]fallypride. We interrogate presynaptic integrity and postsynaptic dopamine receptor availability, and compared these to non-transgenic rats.

*LRRK2* mutant rats displayed similar [^18^F]FDOPA uptake to non-transgenic animals, consistent with intact dopamine synthesis in striatal axons. However, *LRRK2*-R1441C rats demonstrated greater binding of [^18^F]fallypride than *LRRK2*-G2019S or non-transgenic controls, exhibiting regionally selective binding increase in the dorsal striatum. Immunocytochemical labelling post-mortem confirmed a greater density of D2 receptors in *LRRK2*-R1441C than other genotypes restricted to the dorsal striatum, consistent with upregulation of D2-receptors as a compensatory response to the greater dopamine release deficit observed in this genotype.

These results show that [^18^F]fallypride PET imaging is sensitive to dysregulation of dopamine signalling in the *LRRK2*-R1441C rat, detecting upregulation of D2 receptors that parallels observations in early human sporadic PD. Future studies of candidate therapies could exploit this non-invasive approach to assess treatment efficacy.

## Introduction

Parkinson’s disease (PD) is the second most common human neurodegenerative disease worldwide. It is characterized by progressive dysfunction, degeneration and death of dopaminergic neurons in the substantia nigra pars compacta (SNc), which project to the dorsal striatum. Striatal dopamine deficiency is the core deficit underlying the main motor problems of the disease^1,2^. PD is also a multi-system disorder with behavioural, cognitive, and autonomic features, which can precede motor disease by decades^3^. When motor symptoms establish a PD diagnosis, patients have already lost about 60-80% of their dopaminergic neurons^4,5^.

The mechanisms of PD pathogenesis are incompletely understood, hindering the development of neuroprotective therapies^4^. Genetic factors are thought to contribute to perhaps 10% of PD cases^6^. In recent years, new transgenic animal models have been developed based on known PD-associated genes^7–11^ and are aiding discovery of mechanisms contributing to disease progression from early prodromal stages through to late neurodegeneration, as well as providing new models for identifying biomarkers and testing treatments. Several rodent models have been developed based on mutations in the leucine-rich repeat kinase 2 gene (*LRRK2).* LRRK2 mutations are the most common cause of dominantly inherited PD^12^ and account for around 1% of sporadic PD^13^. The G2019S mutation is the most frequent^14^ but, the R1441C mutation induces a more aggressive phenotype^15^. LRRK2 rodent models reproduce age-dependent and progressive motor and cognitive impairment of PD^10,16–19^.

We have previously generated and described Bacterial Artificial Chromosome (BAC) *LRRK2* transgenic rats, expressing either *LRRK2-*R1441C or *LRRK2-*G2019S forms of the human *LRRK2* genomic locus^10,20,21^. These rats exhibit mild age-dependent impairments in dopamine release in dorsal but not ventral striatum, in the absence of any loss of dopaminergic markers (tyrosine hydroxylase [TH], vesicular monoamine transporter 2 [VMAT2] or dopamine transporter [DAT]) in the striatum^10,20^, but paralleled by more dispersed synaptic vesicles and reduced phosphorylated synapsin^20^. In addition, *LRRK2*-R1441C rats display age-dependent reductions in burst firing of dopamine neurons in SNc and have L-DOPA-responsive motor dysfunction. However, no SNc neurodegeneration or abnormal protein accumulation is detectable in these models, suggesting that they represent an early stage of PD and that nigrostriatal dopaminergic dysfunction can precede detectable pathology in early stages of the disease^10^.

Neuroimaging techniques offer a non-invasive strategy to assess dopaminergic dysfunction early in disease, to monitor disease progression, and to assess experimental therapeutic strategies in PD animal models before translation to humans^22^. However, there is little available data to indicate whether PET studies can reveal modifications to nigrostriatal function in new generation, physiological transgenic models of early stages of PD, prior to overt degeneration. PET imaging studies in humans during onset of DA degeneration reveal reductions in dopaminergic presynaptic markers and increased binding of dopamine D2-receptor ligands. Moreover, increased D2-receptor binding in early PD can be localized to the putamen rather than caudate or ventral striatum, and is thought to be an adaptive increase in postsynaptic D2-receptor levels in response to declining levels of dopamine release, rather than due to lower competition for receptor binding from lower levels of endogenous dopamine^23,24^. Increased striatal D2-ligand binding has also been seen in PET imaging in toxin-based animal models of PD, in 6-OHDA rats^25,26^, and MPTP-exposed monkeys^27^ but has not yet been explored or validated in more physiological models of nigrostriatal dysfunction prior to degeneration. One study to date has assessed presynaptic function in *LRRK2-*G2019S rats at 12 months but did not detect significant changes^28^.

Several PET tracers are suitable for imaging dopaminergic function in rodents. The tracer 6-[^18^F]fluoro-L-DOPA ([^18^F]FDOPA) is a marker of pre-synaptic dopamine synthesis capacity, and [^18^F]fallypride indicates dopamine D2/D3 receptor levels^29,30^. We assessed whether PET imaging of [^18^F]FDOPA and [^18^F]fallypride could identify changes to respectively presynaptic function or dopamine receptor levels, using the Waxholm atlas (Figure 1A), and compared susceptible dorsal versus spared ventral striatum (Figure 1B), in *LRRK2-*R1441C and *LRRK2-*G2019S transgenic rat models of early PD, and assessed corresponding D2-receptor immunolabelling.

**Figure 1:**
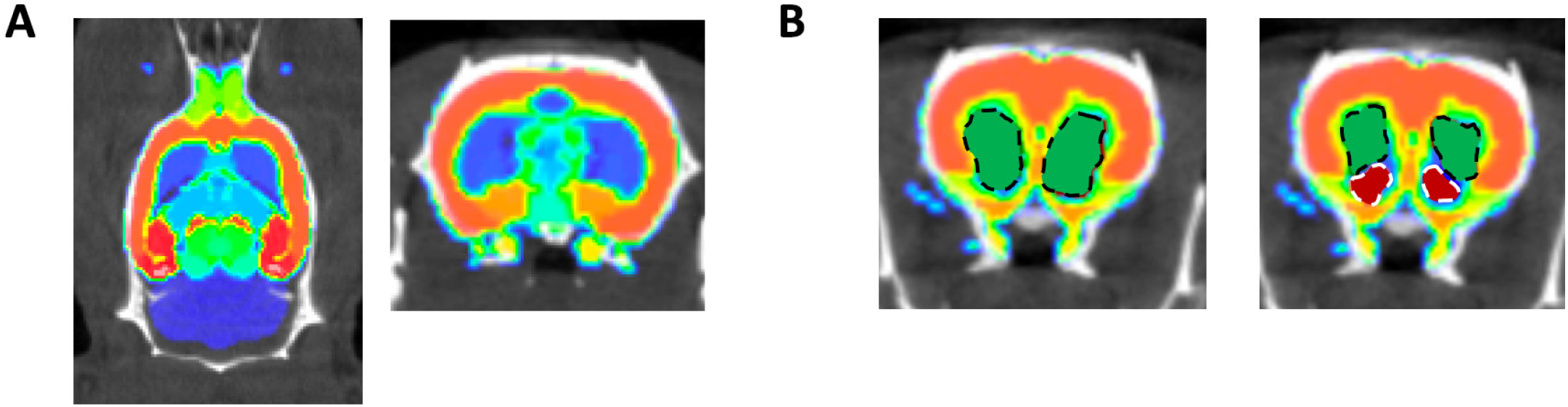
Colour-coded rat brain region templates. **A)** Waxholm rat brain template from^55,56^ superimposed to a representative CT scan from one of the LRRK2 rats included in the PET study. Arbitrary colours to distinguish atlas regions. Left, axial orientation; Right, coronal. **B)** Left, Coronal images showing ROIs calculated using the connected threshold tool and the rat brain template covering left and right striatum (green areas delimited by a black discontinuous line). Right, ROIs calculated for dorsal (green areas delimited by a black discontinuous line) and ventral striatum (red areas delimited by white discontinuous lines).

## Results

### *Dopamine synthesis capacity is preserved in LRRK2*-G2019S *and LRRK2*-R1441C *genotypes*

Specific [^18^F]FDOPA uptake was detected in the striatum, with an acceptable CNR of 3.5 ± 0.93 (Figure 2A). Figure 2B shows an example Patlak plot obtained from the striatum. No significant differences were detected in [^18^F]FDOPA uptake between left and right striata (Table S1). Average uptake data from both striata (Table 2, Figure 2C) indicate that animals carrying either the *LRRK2-*G2019S or the *LRRK2*-R1441C mutation showed no significant deficit in [^18^F]FDOPA uptake in comparison to nTG animals.

**Figure 2:**
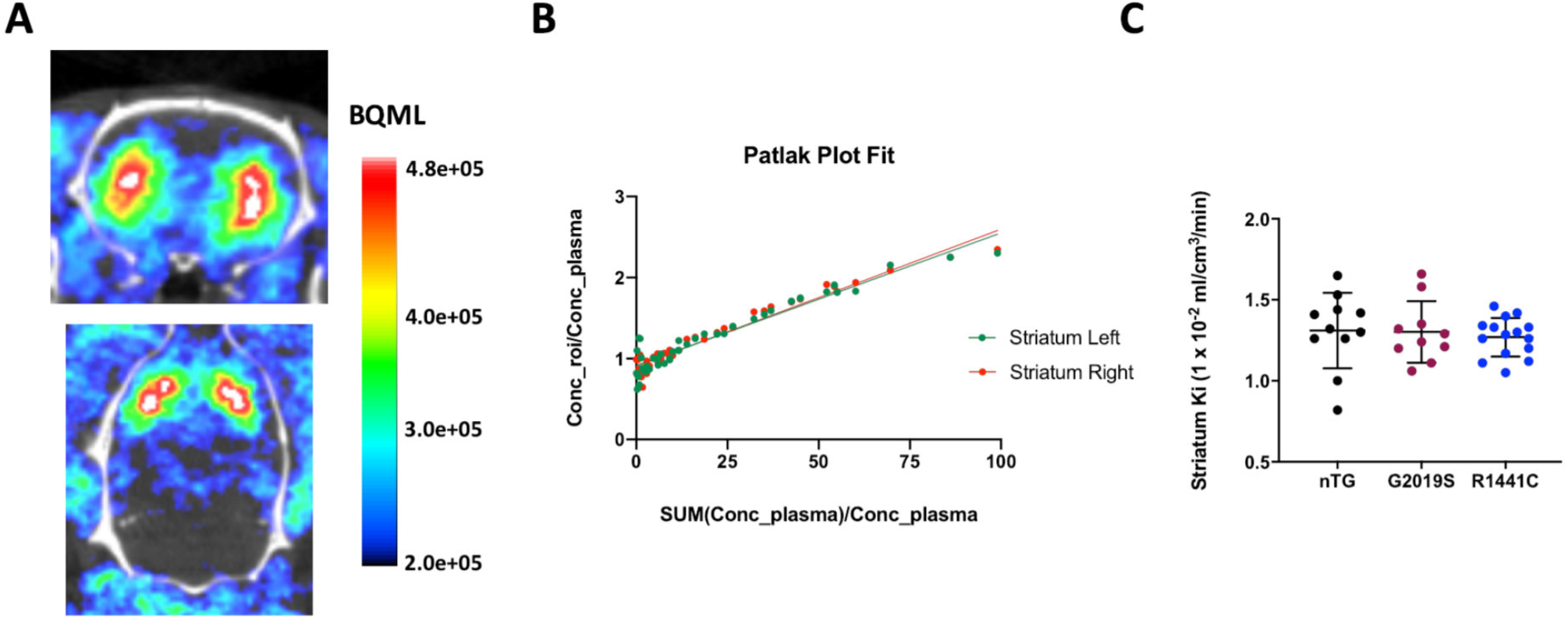
[^18^F]FDOPA PET imaging. **A)** Representative [^18^F]FDOPA PET images acquired in the same animal from coronal (top) and axial (bottom) orientations. The BQML scale range shows the 95% and the 40% of the maximum signal detected as maximum and minimum values respectively. **B)** Patlak plot modelling applied to a representative animal to assess [^18^F]FDOPA dynamic uptake in left and right striatum. **C)** Striatal [^18^F]FDOPA Ki compared between nTG, *LRRK2*-G2019S and *LRRK2*-R1441C genotypes using a One-way ANOVA test followed by pot-hoc comparisons (Tukey HSD). Each dot on the plot represents data from a single rat. * p<0.05.

**TABLE 1:**
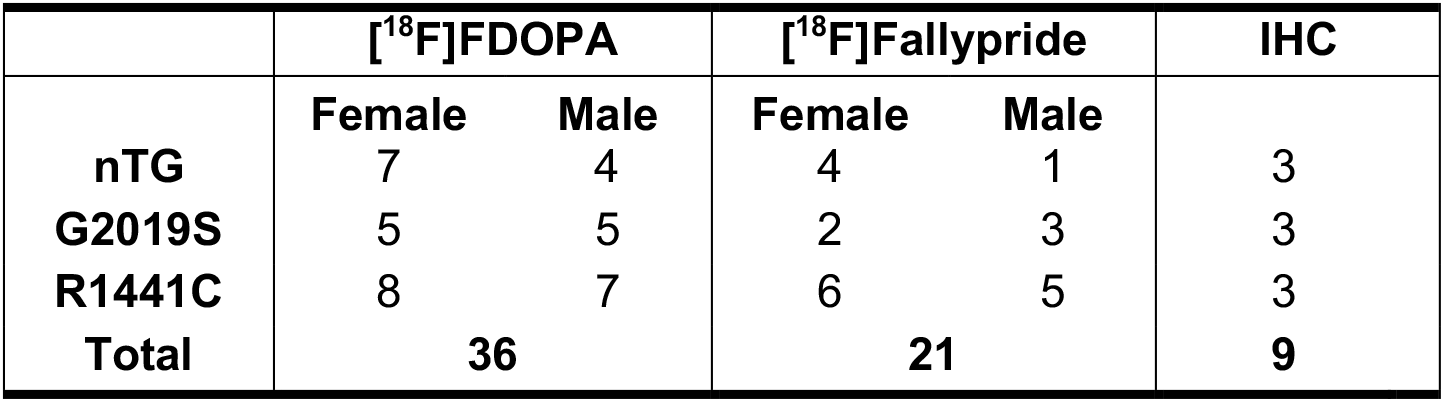
Summary of animal numbers in the study. Numbers of rats by sex, genotype and tracer.

|  | **[<sup>18</sup>F]FDOPA** |  | **[<sup>18</sup>F]Fallypride** |  | **IHC** |
| --- | --- | --- | --- | --- | --- |
|  | **Female** | **Male** | **Female** | **Male** |  |
| **nTG** | 7 | 4 | 4 | 1 | 3 |
| **G2019S** | 5 | 5 | 2 | 3 | 3 |
| **R1441C** | 8 | 7 | 6 | 5 | 3 |
| **Total** | **36** |  | **21** |  | **9** |

**TABLE 2:** [^18^F]FDOPA and [^18^F]fallypride PET tracer dynamic uptake. Average dynamic tracer uptake in the striatum of animals in the 3 groups investigated. [^18^F]fallypride DVR in dorsal and ventral striatal segmentations is also shown. Data represent mean ± SD. One- or Two-way ANOVA tests followed by post-hoc comparisons (Tukey HSD) were used to compare between genotypes (*p < 0.05; **p < 0.01) and between regions within genotype: ^§^p < 0.01; ^§§^p < 0.001.

|  | **[<sup>18</sup>F]FDOPA Ki**<br>**(x10<sup>-2</sup> ml/cm<sup>3</sup>/min)** | **[<sup>18</sup>F]fallypride DVR** |  |  |
| --- | --- | --- | --- | --- |
|  |  | **Total striatum** | **Dorsal** | **Ventral** |
| nTG | 1.31 ± 0.23 | 15.99 ± 4.05 | 15.92 ± 3.96<sup>§</sup> | 9.96 ± 1.92 |
| G2019S | 1.30 ± 0.19 | 16.28 ± 3.15 | 15.97 ± 2.53<sup>§</sup> | 9.73 ± 0.76 |
| R1441C | 1.27 ± 0.12 | 20.93 ± 2.19* | 21.21 ± 2.46<sup>§§**</sup> | 10.55 ± 1.16 |

### D2/D3 receptor binding is upregulated in the dorsal striatum in aged LRRK2-R1441C rats

D2/D3-receptor ligand [^18^F]fallypride PET imaging of resulted in a striatal CNR of 19.6 ± 3.88, giving clear delineation of the rat striatum compared to background (Figure 3A). Logan Reference Plots of tracer binding in an example animal are shown in Figure 3B. No tracer uptake difference was found between left and right striata (Table S3).

**Figure 3:**
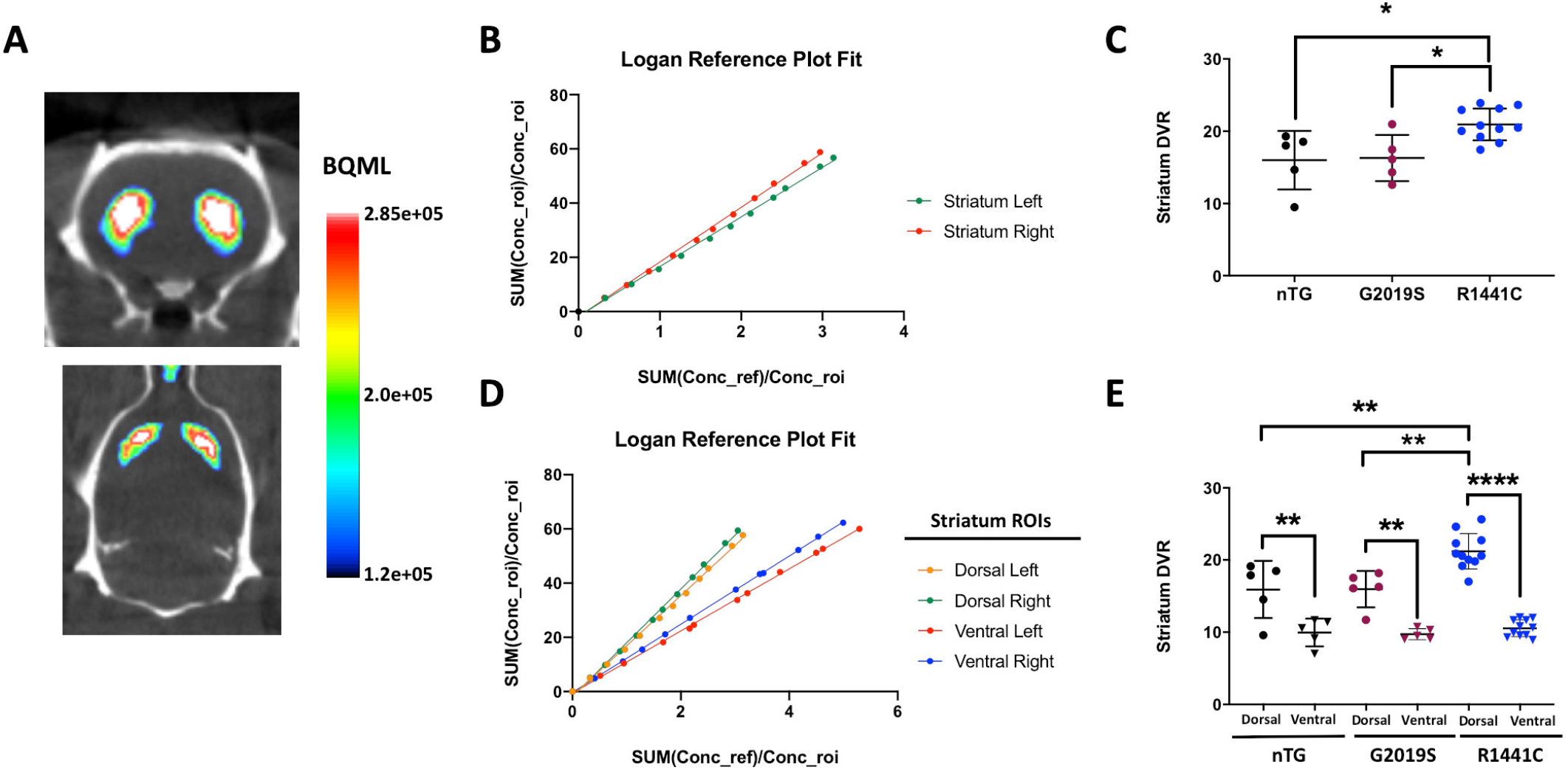
[^18^F]fallypride PET imaging reveals elevated levels in dorsal striatum of *LRRK2*-R1441C rats. **A)** Representative [^18^F]fallypride PET images acquired in the same animal from coronal (top) and axial (bottom) orientations. The BQML scale range shows the 95% and the 40% of the maximum signal detected as maximum and minimum values respectively. **B)** Logan Reference plot modelling applied to the representative case to assess [^18^F]fallypride dynamic uptake in left and right striatum (One-way ANOVA and Tukey). **C)** Striatal [^18^F]fallypride DVR compared between nTG, *LRRK2*-G2019S and *LRRK2*-R1441C groups. **D** and **E** show the same analysis approach described in B and C but segmenting the striatum in dorsal and ventral regions Two-way ANOVA for genotype and region, post-hoc Tukey tests: *p<0.05; **p<0.01; ****p<0.0001.

The level of whole striatal [^18^F]fallypride varied with genotype (One-Way ANOVA, F_2,18_ = 0.422, p=0.006), with *LRRK2-R1441C* rats showing significantly higher binding compared to nTG and *LRRK2-G2019S* groups (Table 2, Figure 3C; post-hoc Tukey tests, p = 0.015 and p = 0.023 respectively). Comparison of the ventral and dorsal subdivisions of striatum, revealed significant differences between genotypes and regions, and a significant interaction (Two-way ANOVA, F_2,36_ = 5.50, p = 0.008). There was a significant difference between dorsal and ventral striatal [^18^F]fallypride binding (Two-way ANOVA, F_1,36_ = 106.9, p < 0.001), owing to greater [^18^F]fallypride binding in dorsal striatum compared to ventral striatum in each of the three genotypes investigated (nTG, p = 0.002; G2019S, p = 0.002 and R1441C, p < 0.001) (Table 2, Figures 3D,E). Furthermore, the significant effect of genotype (F_2,36_ = 9.38, p = 0.001), resulted from [^18^F]fallypride binding within the dorsal striatum of *LRRK2*-R1441C rats being significantly greater than in nTG (p = 0.001) and *LRRK2*-G2019S (p = 0.001) rats. There was no difference in [^18^F]fallypride binding between genotypes for the ventral striatum.

### D2 receptor immunoreactivity is elevated in dorsal striatum in LRRK2-R1441C rats

[^18^F]fallypride binds D2 and D3 receptors, but D3 receptor levels are very low in the striatum in comparison to those of D1 and D2 receptors^31^. To identify whether [^18^F]fallypride binding corresponded to D2-receptor levels, D2 receptor expression in the 3 genotypes (nTG, *LRRK2*-G2019S and *LRRK2*-R1441C) was assessed using indirect immunofluorescence (Figure 4A) in dorsal and ventral striatum. There were significant differences in density of immunolabelling between genotypes and regions, and a significant interaction (Two-way ANOVA, F_2,12_ = 7.08, p = 0.009). There was a significant difference between dorsal and ventral striatal immunolabelling (Two-way ANOVA, F_1,12_ = 25.38, p<0.001), owing to greater D2-receptor density in dorsal striatum compared to ventral striatum in *LRRK2*-R1441C rats only (Tukey post-hoc test, p=0.001). The significant effect of genotype (F_2,12_ = 9.38, p=0.004) was due to greater D2-receptor levels in dorsal striatum of *LRRK2*-R1441C rats than in any other region of any other genotype (Figure 4B).

**Figure 4:**
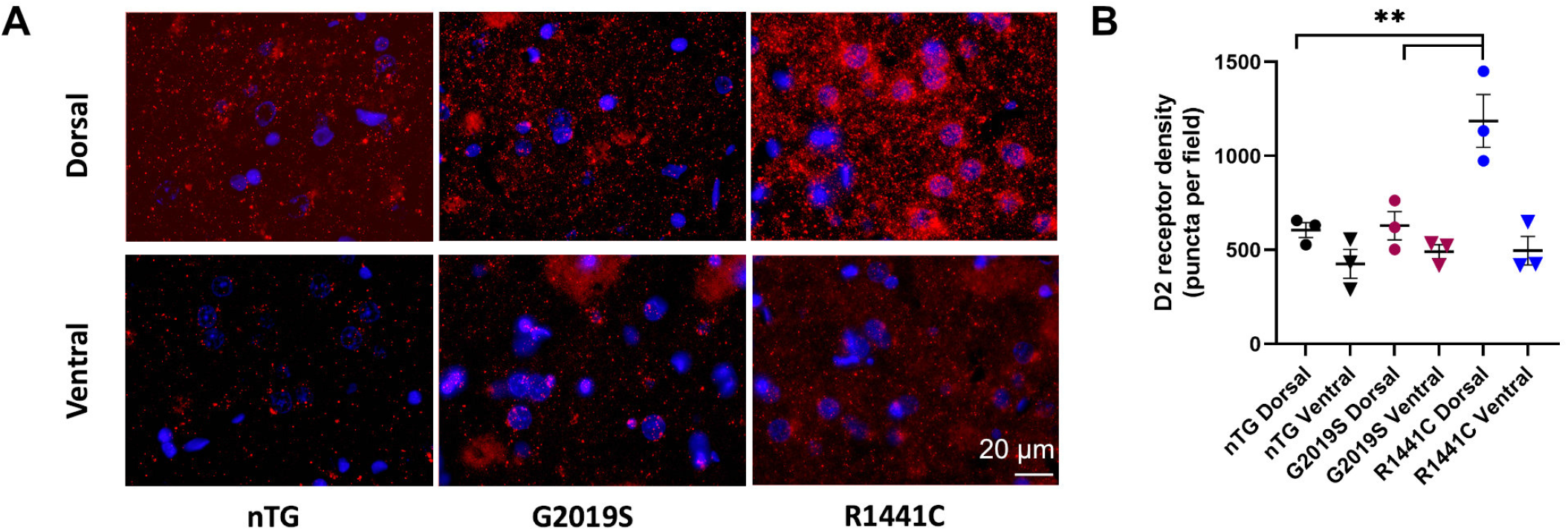
D2-receptor density is elevated in dorsal striatum of *LRRK2*-R1441C rats. **A)** D2 receptors visualised with immunofluorescence staining. Scale bar 20 µm. **B)** Quantification of D2-receptor density, n=3 per genotype. Data represent mean ± SEM. Post-hoc Tukey tests, for clarity, only comparisons between genotypes for dorsal striatum are indicated **p<0.01

## Discussion

We used non-invasive PET imaging to test the utility of the imaging technique for assessing nigrostriatal dysfunction in early-stage PD models of wild-type and mutant human LRRK2-expressing rats. [18F]fallypride imaging identified upregulation of dopamine receptor binding in the dorsal striatum of LRRK2-R1441C transgenic rats, which corresponded to the upregulation of D2-receptor density identified with immunolabeling. Dysregulation of dopamine release and dopamine neuron physiology had previously been identified in this model^10^, and we speculate that relative dopamine depletion at the post-synaptic neuron drives upregulation of D2 receptor density. This is similar to observations in a lesional macaque model^32^ and to PET studies in early human PD^23,24,33^. Another group found upregulation of striatal D2 receptors in an alpha-synuclein overexpressing PD rat model^34^, suggesting that this feature is shared with our chosen model of PD.

*LRRK2* BAC transgenic rats have been characterised in several measures of presynaptic and behavioural function^10,20,21^. Both G2019S and R1441C mutations induce dysregulation of the endocytic pathway and changes in the distribution of synaptic vesicles in dopaminergic neurons. Animals carrying either mutation develop motor and cognitive impairments at an advanced age and display deficits in evoked dopamine release in dorsal striatum, but not in ventral striatum, compared to nTG rats. Dopamine release deficits are greater in *LRRK2*-R1441C than *LRRK2*-G2019S rats. In addition, *LRRK2*-R1441C rats only show a lower frequency of bursts and a lower percentage of spikes fired in bursts compared to nTG rats, as well as a more severe disease phenotype^10^. However, post-synaptic dopamine receptor levels and function had not previously been assessed in these models.

[^18^F]fallypride is a high affinity and highly selective dopamine D2/D3 receptor ligand with negligible affinity for other neuroreceptors^35^, giving it favourable characteristics for imaging dopamine receptor levels in this PD rat model. Its performance has been characterised in rodents^36,37^, including toxin-induced models of PD^26^, but not previously in transgenic models of early PD. We found greater dopamine receptor binding in the *LRRK2-R1441C* genotype than in the other genotypes, which was regionally selective to the dorsal striatum. The dopamine receptor upregulation found in *LRRK2*-R1441C rats is complementary to the impairments in evoked dopamine release we saw previously in this model^10^: These impairments, as measured by sub-second fast scan cyclic voltammetry, are observed in the dorsal, but not the ventral striatum in *LRRK2* mutant aged rats, with a more severe deficit in *LRRK2-*R1441C than *LRRK2*-G2019S rats. Dopamine receptor upregulation in the *LRRK2-*R1441C rat dorsal striatum is consistent with early human PD where, similar to our data, an increase in dopamine receptor density is seen in the putamen, but not the caudate nucleus^24^. In human PD, initial upregulation of dopamine receptors is thought to be a post-synaptic response to dopamine input loss. It is reversed in advanced PD, where a decrease of dopamine receptor density is seen^38–40^. Dopamine receptor upregulation is a candidate compensation mechanism that allows the human motor system to function until the striate dopamine depletion is advanced^41,42^. Now, it also provides a readout of early dopaminergic dysfunction, and we have shown that this can be detected by [^18^F]fallypride PET in the *LRRK2*-R1441C model.

Differences in dopamine receptor tracer binding can reflect dopamine receptor availability^26^ resulting from either lower dopamine occupancy from low release levels^43,44^, or from greater receptor levels^45^ or both. We validated here that differences in [18F]fallypride binding between regions and genotypes reflected differences in the density of the D2 receptor. The greater [18F]fallypride binding and D2-receptor levels in *LRRK2-*R1441C compared to *LRRK2-* G2019S rats is consistent with greater upregulation of post-synaptic D2 receptors arising from the more severe impairment in dopamine release seen in the *LRRK2*-R1441C rat. The more severe phenotype seen with the R1441C rather than the G2019S mutation might in part arise from the fact that these mutations reside in different enzymatic domains of the *LRRK2* gene (the GTPase versus kinase domains respectively), and therefore exert differing effects on the enzymatic activity of *LRRK2*, and consequentially, differing impact on cellular function^21^. It cannot arise from simple concentration differences, as the expression of mutant LRRK2 in the G2019S genotype is greater than in the R1441C model^10^.

The *LRRK2* BAC transgenic models selected for this study exhibit age-dependent and L-DOPA responsive motor dysfunction. In the absence of widespread cell death in this model^10^, the L-DOPA-rescuable PD phenotype suggests that axonal (synaptic) dysfunction precedes overt degeneration of the dopaminergic neurons^46^, causing functional dopamine “deprivation” of postsynaptic neurons. PD animal models capturing α-synuclein pathology have also reported early changes in striatal dopamine release without concomitant differences in striatal dopamine content^8,47^, supporting the hypothesis that synaptic dysfunction precede overt neurodegeneration. Our findings here reveal that the synaptic adaptations occurring without frank dopamine neuron degeneration extend to an upregulation in striatal dopamine receptor density in dorsal striatum, that can be detected with a non-invasive approach of PET imaging.

In conclusion, [^18^F]fallypride PET imaging is sensitive to early dopaminergic dysfunction in the striatum in the LRRK2-R1441C rat model of PD. This non-invasive approach could be used in the future to assess efficacy of new candidate therapies longitudinally in the model.

## Material and Methods

### Animals

*LRRK2* BAC transgenic rats on a Sprague-Dawley background were generated and genotyped as previously reported^10^. Forty-one aged animals (age 21-22 months) including both males and females (weight 350-945 g) were used in this study: non-transgenic rats (nTG, n=11), and two mutant lines, expressing either the *LRRK2*-G2019S (n=10) or *LRRK2-*R1441C (n=15) mutant forms of the human gene (Table 1).

All experiments were conducted in accordance with the Animals (Scientific Procedures) Act, 1986 (United Kingdom) and the University of Oxford Policy on the Use of Animals in Scientific Research, and the ARRIVE guidelines^48^. As previously described, LRRK2-expressing BAC transgenic rats were housed with littermate controls^10^. Rats were housed in a 12 h light-dark cycle with ad-libitum access to food and water. Both sexes were used throughout this study.

### PET acquisition

PET molecular imaging studies were carried out on isoflurane-anaesthetised rats using a Mediso 122S nanoScan PET/CT scanner using an integral multicell XXL rat bed (Mediso Medical Imaging Systems, Budapest, Hungary). 36 animals were scanned with [^18^F]FDOPA, and after complete ^18^F decay, 21 of them underwent an additional scan with [^18^F]fallypride on a different day.

PET data were acquired in list mode into a 128×128×95-matrix. The coincidence window width was set at 6 nsec. PET images were reconstructed with the scanner manufacturer’s nucline software (400 µm resolution, CT based attenuation correction, Tera-tomo 3D reconstruction algorithm, using a full detector model, 4 iterations and 6 subsets). At the end of the PET scan, a 3 minute CT was acquired (50 kVp, 360 rotations) for anatomical localisation and attenuation correction.

General anaesthetic was delivered through a nose cone and maintained during the scanning protocol (2-2.5% isoflurane / 100% oxygen, 1.5 L/min). Temperature was maintained at 37 °C via hot air flow through the bed, and breathing was monitored via a pneumatic sensing pad (∼60 breaths/min). [^18^F]FDOPA and [^18^F]fallypride were produced in house at the Wales Research and Diagnostic PET Imaging Centre (PETIC) on a Trasis all-in-one system via adaptations of literature methods^49,50^. The tail vein was cannulated 15 minutes before scanning using a 22G/25mm cannula (Millpledge MP06222, UK) attached to an Anisite membrane locking cap (Millpledge HB07510, UK).

For [^18^F]FDOPA experiments, animals were injected i.p. with the COMT inhibitor entacapone (40 mg/kg, Glentham Life Sciences GP9755, UK) in a DMSO/saline solution (60/40 v/v, 11.85 mg/ml) adjusted to pH 7 with NaOH solution (1 M, 1-10 µL), 90 minutes before tracer administration. The peripheral Aromatic L-Amino Acid Decarboxylase (AADC) inhibitor benserazide hydrochloride (10 mg/kg, Glentham Life Sciences GP7664, UK) was administered in aqueous solution (10 mg/ml), 60 minutes before tracer injection^51^. Benserazide does not cross the blood-brain barrier. This inhibitor combination is licensed for oral use in PD patients receiving L-DOPA treatment^52^, and ensures that L-DOPA (and here, [^18^F]FDOPA) reaches the brain intact.

[^18^F]FDOPA was injected as a single bolus (151 MBq/kg) followed by saline flush (50 µL). A 90 minutes single bed position PET scan was acquired in 3D list mode starting immediately at the end of radiotracer injection^53^. The raw PET data were sorted into 41 frame-dynamic sinograms as follows: 6 × 10 s, 6 × 30 s, 11 × 60 s, 15 × 180 s, 3 × 600 s.

[^18^F]fallypride scans were carried out with the same scanner set-up and animal preparation used for [^18^F]DOPA, with the exception that no inhibitors are required before tracer injection. [^18^F]fallypride dose was 71 MBq/kg followed by a saline flush (50 µL). A 60 minute PET scan was acquired in 3D list mode starting at 60 minutes post radiotracer injection. Data were sorted into 12x300 s timeframes^44^.

Tracer cold masses injected are shown in Table S2 in the supplementary material.

### Molecular imaging analysis

PET images were analysed using VivoQuant® software with an additional pharmacokinetic modelling module (Invicro, Boston, MA, US). Regions of interest (ROIs) corresponding to left and right striatum were created for every scan using the connected threshold tool. A representative cerebellum region was drawn with the free hand tool, to provide a reference input function for tracer uptake analysis.

Data were processed using two different approaches: i) dynamic analysis of tracer uptake during the full scanning time with multiple-time graphical analysis (Patlak and Logan plots as appropriate), and, for comparison, ii) calculation of tracer uptake from static imaging (standardized uptake value (SUV)) derived from the last 30 minutes of acquisition. Static images were also used to calculate contrast-to-noise (CNR) for both tracers.

[^18^F]FDOPA data were analysed with Patlak plots, analogous to previous approaches^53^. The 10 initial minutes from each scan were discarded for analysis purposes^54^. Left and right target striatal ROI influx estimates (Ki) were averaged and used for group-wise analysis.

[^18^F]fallypride PET imaging was used to assess D2/D3 receptor levels in the striatum of nTG, *LRRK2*-G2019S and *LRRK2*-R1441C animals. Logan reference plots over the entire 60 minute scan were applied to [^18^F]fallypride data. Left and right ROI distribution volume ratio (DVR) values were averaged for each rat and subjected to statistical analysis. The high CNR of [^18^F]fallypride scans allowed further subdivision of the rat striatum into dorsal and ventral sections (Figure 1B). This was not possible in the case of [^18^F]FDOPA as the initially lower CNR was severely decreased after region segmentation, affecting Patlak plot fitting (Supplementary Material, Figure S1). The Waxholm atlas^55,56^ was used in combination with VivoQuant® software for ROI delineation and [^18^F]fallypride DVR was calculated in these sub-regions as described above. The anatomic distribution of the ROIs used in the segmented analysis is shown in Figure S2.

### Immunolabeling

An additional rat cohort (n=9) was used for the immunofluorescence validation of D2 receptor expression levels. Animals were culled under terminal anaesthesia using pentobarbital (0.7 g/kg, i.p.) and transcardially perfused with 0.01 M phosphate buffered saline (pH 7.4) followed by 4% w/v paraformaldehyde in 0.01 M phosphate buffered saline. Paraffin embedded brain sections, cut to a thickness of 8 μm, were dewaxed and rehydrated through a series of alcohol. Slides were blocked in PBS-Tween (0.4% v/v tween) containing 10% v/v normal donkey serum for 60 minutes at room temperature, then incubated with the primary D2 receptor antibody (ab85367, Abcam, UK) (1:250 dilution in PBS-Tween with 10% normal donkey serum) over night at 4°C. Samples were then washed and incubated with the secondary antibody (A-21207, ThermoFisher) (1:1000 dilution in PBS-Tween) for 1 hour at room temperature, thereafter, washed and incubated in 1 μg/ml DAPI (Life Technologies). Coverslips were mounted onto Superfrost plus (VWR) microscope slides using FluorSave reagent (Millipore). Samples were imaged using an Invitrogen EVOSTM FL Auto cell imaging system at 60x magnification. Eight images were acquired in the dorsal and 6 in the ventral striatum for each animal. D2 receptor staining was quantified with the ImageJ software (v1.52n, NIH, USA) using the particle analysis module after applying background subtraction (rolling:35) and the unsharp masking filter (radius:1; mask weight:0.8).

### Statistical analysis

Statistical analyses were performed using SPSS 23 (IBM, USA) and GraphPad Prism 8 and 9 (GraphPad Software, USA). Data were analysed by One- or Two-way analysis of variance (ANOVA), or Student’s t-tests. In instances when group size did not allow for standard tests of normality (i.e. Shapiro-Wilk, Kolmogorov-Smirnov), the normality of the residuals was assessed instead. Post-hoc tests following ANOVAs were conducted using Tukey HSD correction. Two-tailed levels of significance were used and p < 0.05 was considered statistically significant. Data are expressed as mean ± standard deviation (SD) throughout the manuscript.

## Acknowledgments

This work was supported by the Oxford Parkinson’s Disease Centre, funded by the Monument Trust Discovery Award from Parkinson’s UK (Grant J-1403), with the support of the National Institute for Health Research (NIHR) Biomedical Research Centres based at Oxford University Hospitals NHS Trust, Oxford Health NHS Foundation Trust and University of Oxford, and the NIHR Clinical Research Network. Dr Klein acknowledges support from the NIHR Oxford Health Clinical Research Facility. P.J.M. was supported by the Medical Research Council UK (award MC_UU_12024/2). The Wellcome Centre for Integrative Neuroimaging is supported by core funding from the Wellcome Trust (203139/Z/16/Z).

## Author contribution statement

J.C.K., C.E.M., R.W-M, T.D-G., N.C-R., P.J.M., and S.J.C. contributed to the study conception and design. Material preparation, data collection and analysis were performed by T.D-G., S.P., N.C-R., M.C., E.L.L., C.M, D.H. and J.M. The first draft of the manuscript was written by T.D-G, N.C-R and J.C.K. and all authors commented on following versions of the manuscript. All authors read and approved the final manuscript.

## Compliance with Ethical Standards

### Ethical approval

All applicable international, national, and/or institutional guidelines for the care and use of animals were followed. All procedures performed in studies involving animals were in accordance with the ethical standards of the institution at which the studies were conducted.

All animal procedures were carried out under the United Kingdom Animals (Scientific Procedures) Act (1986).

