## Supplementary Tables and Figures for "Dopamine D2 receptor upregulation in dorsal striatum in the *LRRK2*-R1441C rat model of early Parkinson’s disease revealed by *in vivo* PET imaging"

**TABLE S1:** Average dynamic [$^{18}$F]FDOPA uptake in the left and right striatum of animals in the 3 groups investigated. Data represent Mean ± SD. Paired t-Tests used for statistical comparisons.

| | [$^{18}$F]FDOPA Ki (1 x $10^{-2}$ mL/cm3/min) | | |
|---|---|---|---|
| | **Left Striatum** | **Right Striatum** | **p value** |
| **nTG** | 1.3 ± 0.22 | 1.3 ± 0.24 | 0.361 |
| **G2019S** | 1.3 ± 0.18 | 1.3 ± 0.21 | 0.537 |
| **R1441C** | 1.3 ± 0.11 | 1.3 ± 0.13 | 0.322 |

**TABLE S2:** Average cold tracer amount (mM) injected per animal in each group. Data represent Mean ± SD. Kruskal-Wallis test used for statistical comparisons.

| | **Average cold tracer injected per animal (mM)** | |
|---|---|---|
| | **DOPA** | **Fallypride** |
| **nTG** | 1.6E-03 ± 3.49E-04 | 6.1E-04 ± 9.58E-05 |
| **G2019S** | 1.5E-03 ± 3.47E-04 | 6.5E-04 ± 1.50E-05 |
| **R1441C** | 1.6E-03 ± 3.24E-04 | 5.1E-04 ± 2.08E-04 |
| **p value** | 0.61 | 0.88 |

**TABLE S3:** Average dynamic [$^{18}$F]Fallypride uptake in the left and right striatum of animals in the 3 groups investigated. Data represent Mean ± SD. Paired t-Tests used for statistical comparisons.

| | [$^{18}$F]Fallypride DVR | | |
|---|---|---|---|
| | **Left Striatum** | **Right Striatum** | **p value** |
| **nTG** | 15.9 ± 4.02 | 16.0 ± 4.08 | 0.853 |
| **G2019S** | 16.4 ± 3.24 | 16.2 ± 3.13 | 0.189 |
| **R1441C** | 20.8 ± 2.51 | 21.1 ± 1.93 | 0.263 |

### SUPPLEMENTARY FIGURES

**FIGURE S1:** Patlak plot modelling applied to a representative case (1 rat) to assess [18F]FDOPA dynamic uptake in **A)** left and right striatum and **B)** Dorsal and ventral left/right striatum. Region segmentation reduces significantly the CNR affecting the model fitting.

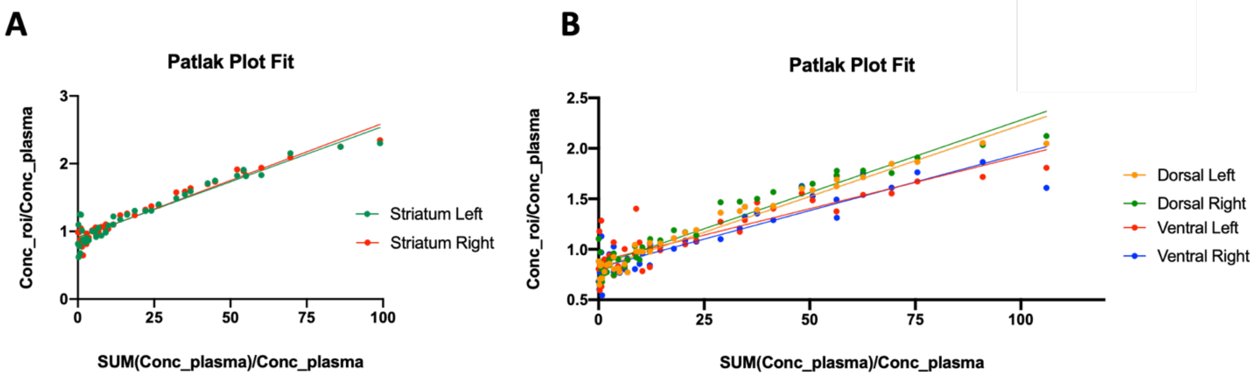

**FIGURE S2:** Anatomical location of the striatum segmentations used for [$^{18}$F]Fallypride analysis shown on Paxinos rat brain atlas (coronal and sagittal orientations). Dorsolateral striatum regions are delimited by green lines and ventral striatum regions (nucleus accumbens) are delimited by red lines.

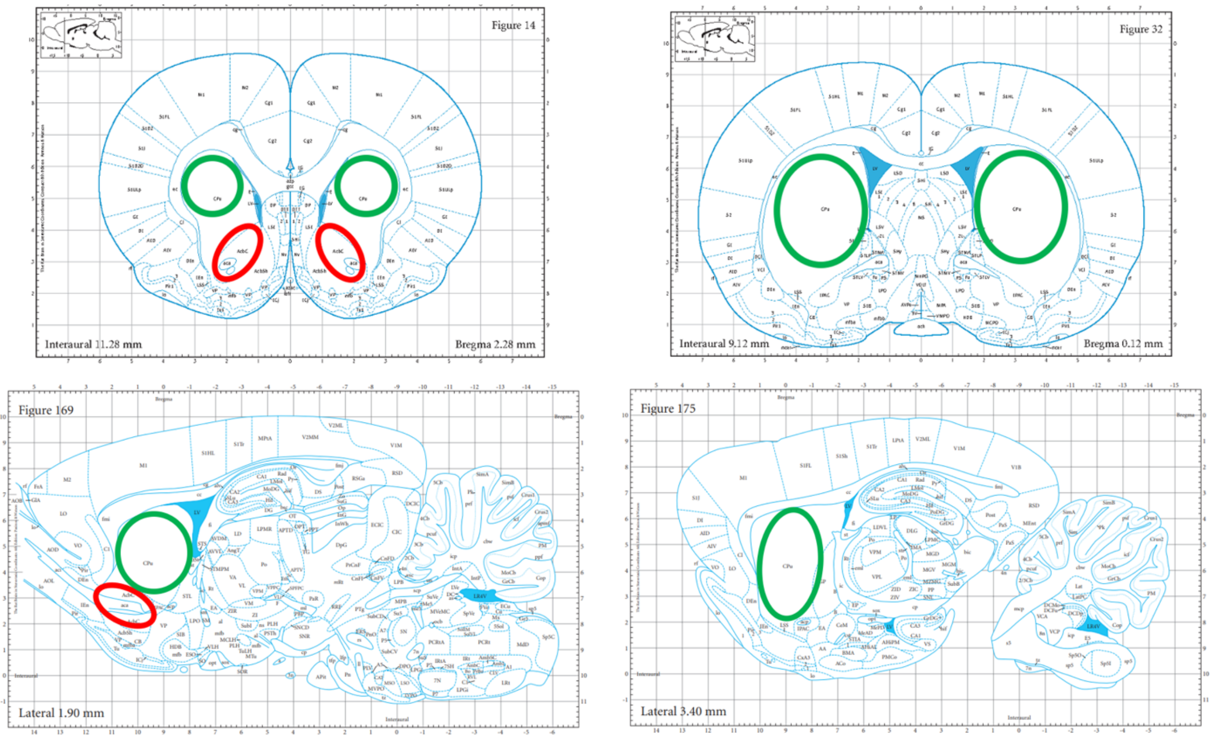
